# Classifying Asian Rice Cultivars (*Oryza sativa* L.) into *Indica* and *Japonica* Using Logistic Regression Model with Publicly Available Phenotypic Data

**DOI:** 10.1101/470351

**Authors:** Bongsong Kim

## Abstract

This article introduces how to implement the logistic regression model (LRM) with phenotypic variables for classifying Asian rice (*Oryza sativa* L.) cultivars into two pivotal subpopulations, *indica* and *japonica*. This study took advantage of publicly available data attached to a previous paper. The classification accuracy was assessed using an area under curve (AUC) of a receiver operating characteristic (ROC) curve. Given 24 phenotypic variables for 280 *indica/japonica* accessions, the LRMs were fitted with up to six phenotypic variables of all possible combinations; the highest AUC accounts for 0.9977, obtained with six variables including panicle number per plant, seed number per panicle, florets per panicle, panicle fertility, straighthead susceptibility and blast resistance. Overall, the more variables there are, the higher the resulting AUCs are. The ultimate purpose of this study is to demonstrate the *indica/japonica* prediction ability of the LRM when applied to unclassified Asian rice cultivars. To estimate the *indica/japonica* prediction accuracy, ten-fold cross-validations were conducted 100 times with the 280 *indica/japonica* accessions using the LRM with parameters that yielded the highest AUC. The resulting prediction accuracy accounted for 0.9779. This suggests that the LRM promises to be a highly effective *indica/japonica* prediction tool using phenotypic variables in Asian cultivated rice.

## Introduction

Asian cultivated rice (*Oryza Sativa* L.) is known to have five subpopulations which are *indica, temperate japonica, tropical japonica, aus* and *aromatic* (Glaszmann, 1987). Of these, *indica* and *japonica*, comprised of *temperate japonica* and *tropical japonica*, are known as pivotal subpopulations; *aus* is known to be related to *indica* and *aromatic* as a medium type between *indica* and *japonica* (Garris et al, 2005; Zhao et al, 2011; McCouch et al, 2016; Chin et al, 2017). There are genetic barriers particularly between *indica* and *japonica*, which often challenge rice trait improvements by breeding (Chen et al, 2008; Kim et al, 2009; Zhu et al, 2017). Because each subpopulation often has desirable characteristics for cultivars, overcoming the genetic barriers between subpopulations will help rice breeders freely introgress desirable genes originating from different subpopulations into an elite line. This can eventually minimize the breeding costs and maximize the sustainability of rice production being threatened by ongoing climate changes, rapidly increasing human population and rapidly decreasing rice cultivation land. To this end, classification of Asian cultivated rice is considered a fundamental study and has been extensively conducted (Garris et al, 2005; Kim et al, 2009; Zhao et al, 2011; McCouch et al, 2016; Chin et al, 2017). With regard to Asian rice classification, genomic tools have been popular to date because genomic data reflect the variability of the subpopulation characteristics at a molecular level. However, the sole use of genomic data may have limitations to the full understanding of the Asian rice classification because (1) some subpopulation-associated traits are related to cytoplasmic effects (Bao et al, 2002; Zhao et al, 2010; Tao et al, 2011; Dai et al, 2017), and (2) a genomic data set per se is often biased while selecting polymorphic markers (Kim and Beavis, 2017). To overcome the gap that genomic data cannot fill, the use of phenotypic data is reasonable because it comprehends the genomic and cytoplasmic effects.

This article addresses how to use phenotypic data for classifying Asian rice cultivars into *indica* and *japonica*. The analytical method employed is the logistic regression model (LRM) in which a response variable is binary between *indica* and *japonica*, and predictors (phenotypic variables) are quantitative. This study spent no cost on data generation by taking advantage of a publicly available data source (http://www.ricediversity.org) containing information on 413 Asian rice cultivars. The data-related article was previously published (Zhao et al, 2011). This study demonstrates that the LRM is a promising tool that makes highly accurate *indica/japonica* predictions of unclassified Asian rice cultivars by phenotype.

## Materials and Methods

### Phenotypic data

A set of phenotypic data and subpopulation information was used, which was originally generated and analyzed by Zhao et al. (2011). The data set was freely obtained at http://ricediversity.org/data/sets/44kgwas, consisting of 413 accessions originated from 82 countries. Out of 32 phenotypic variables, 24 variables were selected because they are quantitative and not confined to certain geography. The selected variables can be divided into morphology (culm habit, flag leaf length, flag leaf width), yield components (panicle number per plant, plant height, panicle length, primary panicle branch number, seed number per panicle, florets per panicle, panicle fertility), seed morphology (seed length, seed width, seed volume, seed surface area, brown rice seed length, brown rice seed width, brown rice volume, seed length width ratio, brown rice length width ratio), stress tolerance (straighthead suseptability, blast resistance) and quality (amylose content, alkali spreading value, protein content). Every accession in the data set belonged to one of the following five subpopulations: *admixed* (62), *aromatic* (14), *aus* (57), *indica* (87), *temperate japonica* (96) and *tropical japonica* (97). In this study, *temperate japonica* and *tropical japonica* were combined into *japonica*.

### *Indica/japonica* classification based on logistic regression model

Only the *indica* and *japonica* accessions were selected to establish the LRM to perform *indica/japonica* classifications using phenotypic variables. Fitting the LRM generates parameters corresponding to predictors. Applying phenotypic observations from each single accession to the LRM with estimated parameters produces a value restricted between 0 and 1, which classifies the accession into either *indica* or *japonica* at 0.5. Suppose that an nD is a set containing all possible combinations of *n* different predictors, e.g. 1D contains all predictors; 2D contains all possible pairs of predictors. In this study, I established 6 sets from 1D to 6D. The LRM was established with every selection of each set to estimate parameters as follows:

1. Given 1D,

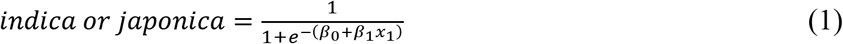
2. Given 2D,

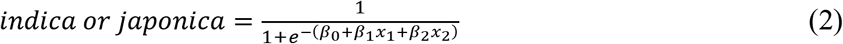
3. Given 3D,

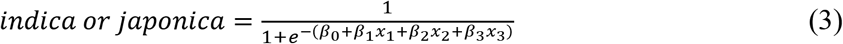
4. Given 4D,

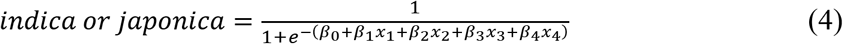
5. Given 5D,

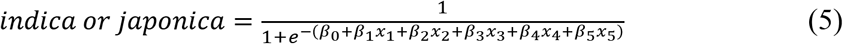
6. Given 6D,

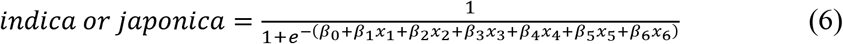

where *indica or japonica* is a value ranging between 0 and 1 which determines *indica* or *japonica* at 0.5; *β*_0_ is the constant term; *β*_1_, *β*_2_, *β*_3_, *β*_4_, *β*_5_ and *β*_6_ are the parameters for the predictors *x*_1_, *x*_2_, *x*_3_, *x*_4_, *x*_5_ and *x*_6_, respectively.

The uses of the 6 sets (1D to 6D) aimed at (1) finding a selection of predictors that yield the best *indica/japonica* classification accuracy and (2) observing the pattern between the number of predictors and classification accuracy. In addition, the LRM with parameters yielding the best classification accuracy was applied to each minor subpopulation (*admixed*, *aromatic*, *aus*) to assign each accession to either *indica* or *japonica* by phenotype.

### Genomic data

The genomic data set consisted of 413 accessions and 36,901 SNPs. This data set was originally generated and analyzed by Zhao *et al*. (2011) and freely obtained at http://ricediversity.org/data/sets/44kgwas. The genomic data set was used to draw dendrograms to classify the rice accessions into the subpopulations.

### *Indica/japonica* classification based on dendrogram

The dendrogram-based *indica/japonica* classifications were performed to compare with the LRM-based *indica/japonica* classifications, for which a genetic distance matrix was calculated, then graphed into a dendrogram. The genetic distances were calculated using the following equation:

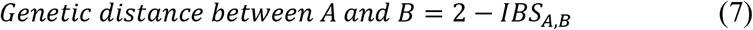

where the *IBS_A,B_* is the IBS (identical by state) coefficient between A and B, which ranges between 0 and 2.

The IBS matrix was calculated using Numericware i (Kim and Beavis, 2017), for which the genomic data set was transformed using Linux Bash. The dendrogram was drawn using R (R Core Team, 2016). In this study, two dendrograms were drawn with (1) only *indica/japonica* accessions and (2) the whole population set.

### *Indica/japonica* classification accuracy

To estimate the *indica-japonica* classification accuracy, a receiver operating characteristic (ROC) curve was implemented using an R package called pROC (Robin et al, 2011). In an ROC space, the *x* and *y* axes ranging between 0 and 1 represent the false positive rate (synonymous to FPR and specialty) and true positive rate (synonymous to TPR and sensitivity), respectively; thus, an area under an ROC curve (AUC) can range between 0 and 1. The closer an AUC is to 1, the higher the *indica-japonica* classification accuracy is.

### *Indica/japonica* prediction accuracy

The LRM can be useful for making *indica/japonica* predictions of unclassified Asian rice cultivars, which is the ultimate purpose of this study. The *indica/japonica* prediction accuracy of the LRM was estimated by averaging values resulting from 100 times iterative ten-fold crossvalidations (CVs).

### R scripts

All of the LRM-associated computations were conducted using R (R Core Team, 2016). The R scripts are freely available at https://github.com/bongsongkim/logit.regression.rice.

## Results

### LRM-based *indica/japonica* classification

The LRM was fitted with each selection in 1D to 6D; the classification accuracy for each selection was estimated into an AUC. Table 1 summarizes the resulting AUCs at each set, which shows that the AUCs tend to increase, as the number of predictors increases. Figure 1 shows the best LRM curve for each set. Across the six sets, the highest AUC accounts for 0.9977, obtained from 6D. The predictors that achieved the maximum AUC in each set are summarized in Table 2; three most frequent predictors are panicle number per plant (6 times), blast resistance (4 times) and straighthead susceptibility (4 times). Parameters that achieved the maximum AUC are - 18.745 for panicle number per plant, 148.256 for seed number per panicle, −154.193 for florets per panicle, −216.319 for panicle fertility, and −1.983 for straighthead susceptibility, 1.223 for blast resistance and the constant term is 312.056. The LRM in which the parameters reside is shown as follows:

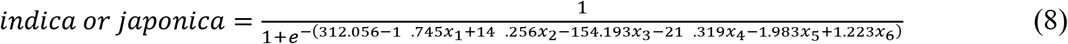

where *indica or japonica* is a value ranging between 0 and 1 which determines *indica* or *japonica* at 0.5;*x*_1_ = the predictor for panicle number per plant; *x*_2_ = the predictor for seed number per panicle; *x*_3_= the predictor for florets per panicle;*x*_4_= the predictor for panicle fertility;*x*_5_= the predictor for straighthead susceptibility; *x*_6_ = the predictor for blast resistance.

**Table 1.** Summary of the resulting AUCs obtained in each set (1D to 6D).

| Set name | Minimum | Median | Mean | Maximum |
| --- | --- | --- | --- | --- |
| 1D | 0.5129 | 0.6937 | 0.6868 | 0.9396 |
| 2D | 0.5192 | 0.7968 | 0.7829 | 0.9762 |
| 3D | 0.5507 | 0.8441 | 0.8413 | 0.9892 |
| 4D | 0.5861 | 0.8789 | 0.8805 | 0.9927 |
| 5D | 0.6101 | 0.9045 | 0.9081 | 0.9968 |
| 6D | 0.6299 | 0.9266 | 0.9283 | 0.9977 |

**Table 2.**
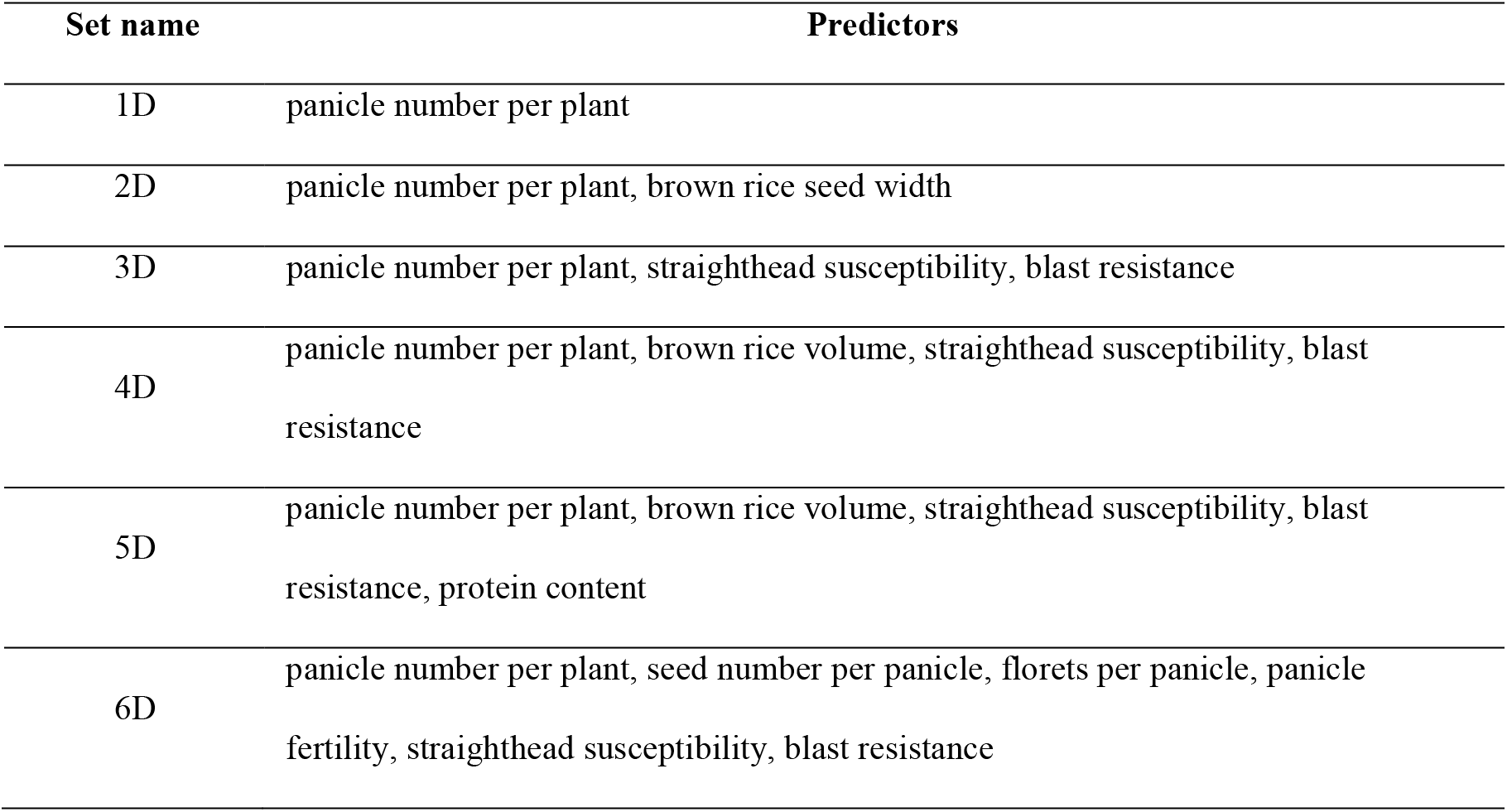
Summary of predictors yielding the best classification accuracy in each set (1D to 6D).

**Figure 1.**
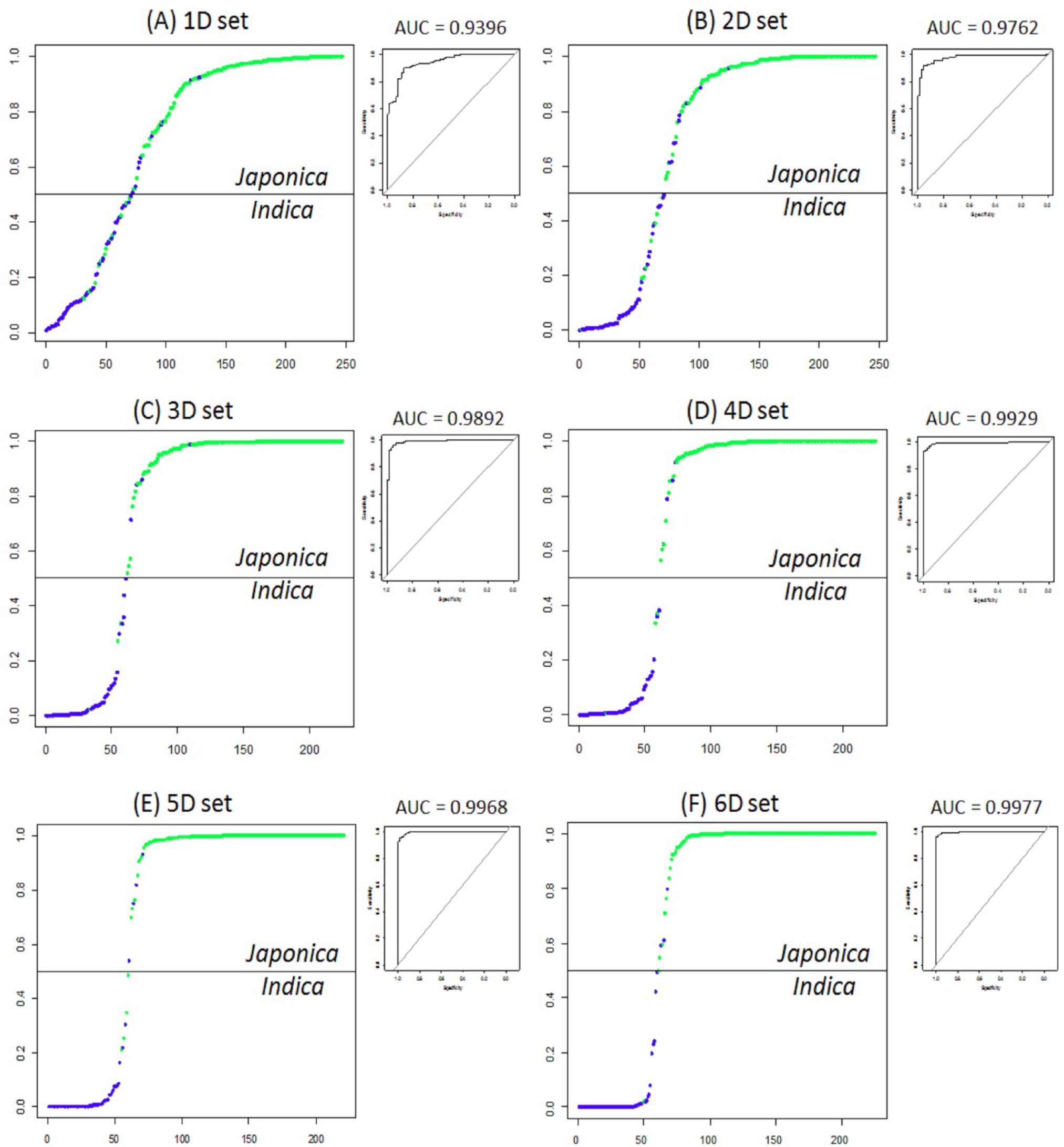
Each panel shows a logistic regression curve graph (left) and an ROC curve graph (upper right) yielding the best classification accuracy in each set (1D to 6D).

### Dendrogram-based *indica/japonica* classification

Figure 2 shows the resulting dendrogram-based *indica/japonica* classification given 280 *indica* and *japonica* accessions, in which the *indica* and *japonica* groups are perfectly divided; and the *japonica* accessions were further perfectly divided into *temperate japonica* and *tropical japonica* (Supplementary Figure 1). This result shows that the dendrogram-based *indica/japonica* classification accuracy accounts for 100 % (AUC = 1).

**Figure 2.**
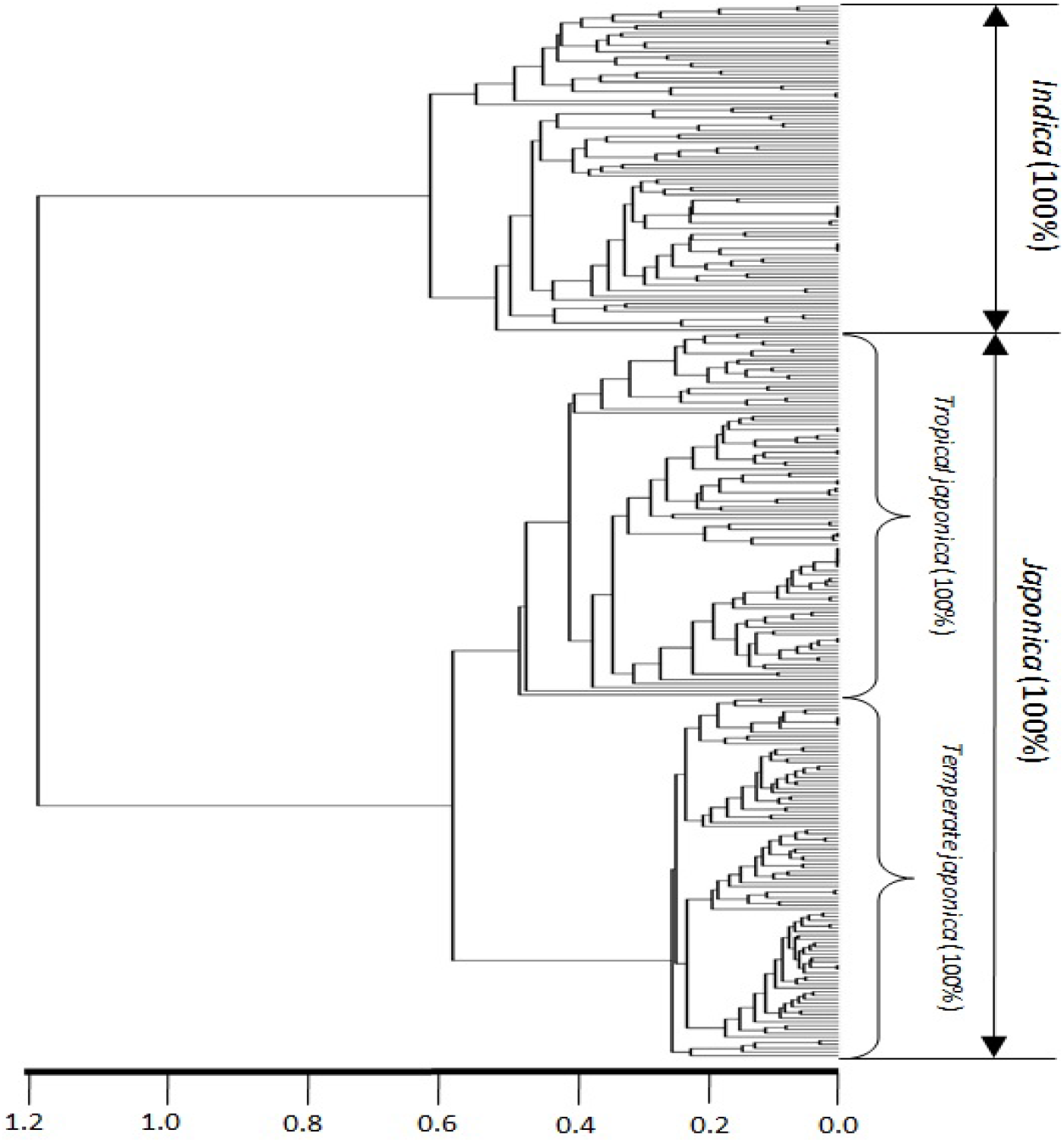
Dendrogram derived from genetic distances between 280 *indica/japonica* accessions using 36,901 SNPs.

### Classifying accessions in each minor subpopulation into *indica* and *japonica*

To investigate how the given accessions in the *admixed, aromatic* and *aus* subpopulations are classified into *indica* and *japonica*, I applied Equation 8 to each subpopulation. Because of missing phenotypic records, the size of each subpopulation was reduced from 62 to 52 for *admixed*, 14 to 12 for *aromatic* and 57 to 26 for *aus*. Figure 3 shows that accessions were assigned to *indica* and *japonica* in ratios of 5:47 for *admixed*, 4:8 for *aromatic* and 20:6 for *aus*, respectively. Meanwhile, the dendrogram drawn with the whole population set (413 accessions) provided two major clades (upper and lower); *temperate japonica, tropical japonica* and *aromatic* formed perfectly separate groups within the upper clade, while *indica* and *aus* formed perfectly separate groups in the lower clade, and the *admixed* accessions were spread across all subpopulations (Supplementary figure 2). Figure 4 shows three Venn diagrams, each of which represents comparison between the LRM-based and dendrogram-based classifications for each minor subpopulation; the agreements account for 92.3 % (48/52) in the *admixed* group, 66.7% (8/12) in the *aromatic* group and 76.9 % (20/26) in the *aus* group.

**Figure 3.**
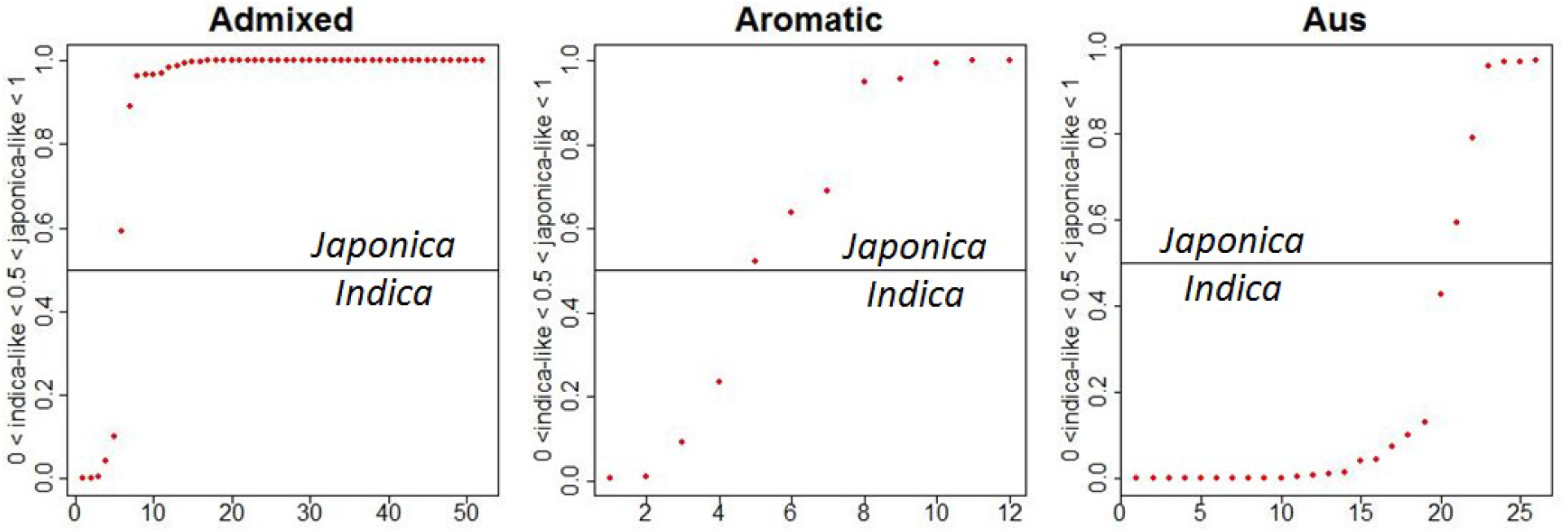
LRM-based *indica/japonica* classification using Equation 8 for each minor subpopulation.

**Figure 4.**
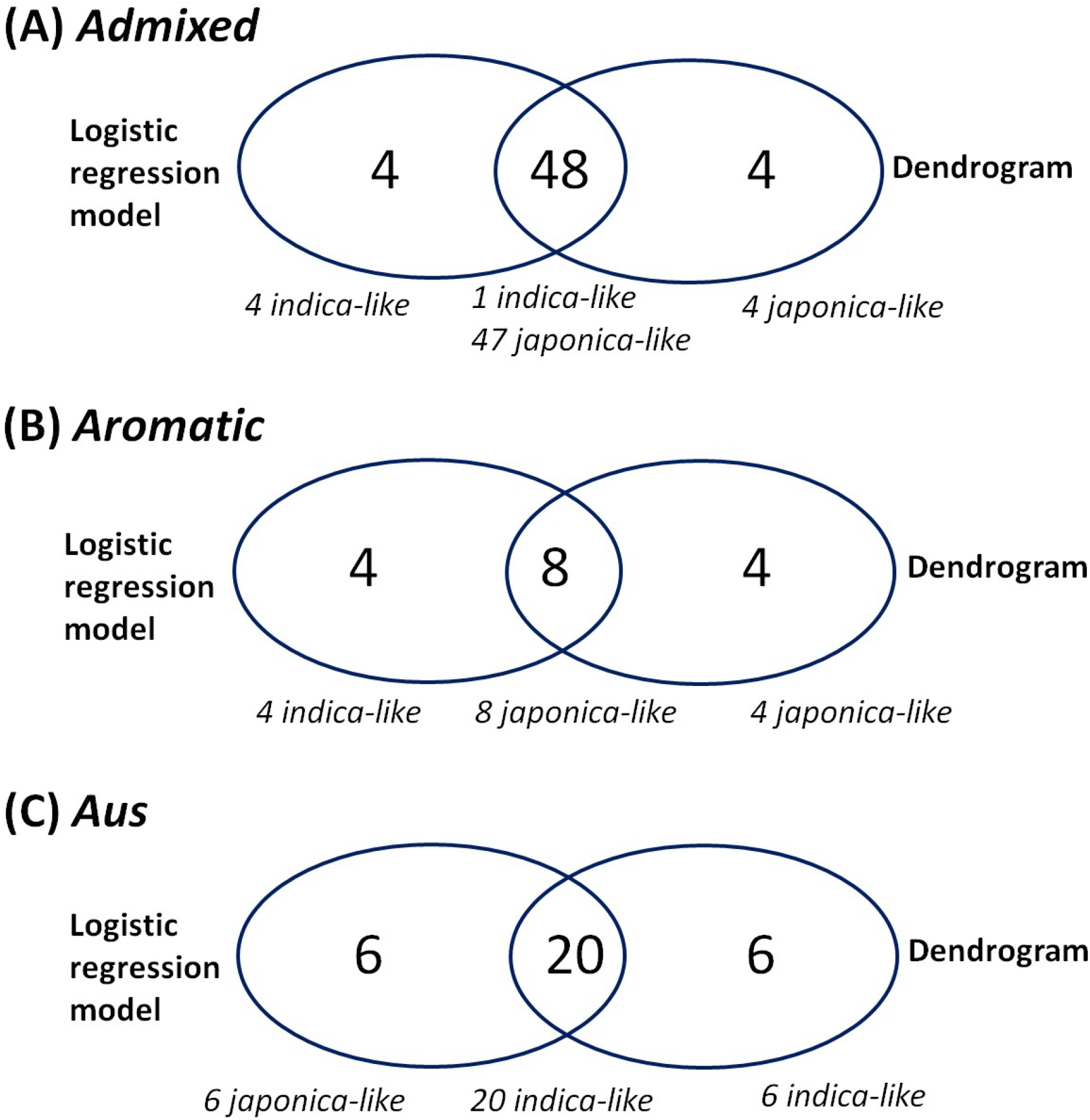
Each panel shows a Venn diagram representing comparison between the LRM-based classification using Equation 8 and the dendrogram-based classification for each minor subpopulation.

### LRM-based *indica/japonica* prediction

The ultimate purpose of this study is to demonstrate the effectiveness of the LRM-based *indica/japonica* predictions when applied to unclassified Asian rice cultivars. To this end, the ten-fold cross-validations using Equation 8 with the 280 *indica/japonica* accessions were repeated 100 times, which resulted in a prediction accuracy of 0.9779. The dendrogram-based classifications cannot fulfill the *indica/japonica* predictions because it is not a predictive method that generates parameters but an estimative method that makes a confirmative classification.

## Discussion

This article demonstrates that the LRM is a powerful tool for classifying Asian rice cultivars into *indica* and *japonica* using phenotypic predictors. AUCs obtained by the LRM gradually increases, as the number of predictors increases (Table 1 and Figure 1). This implies that the variability in a single trait is narrowly distinguishable between *indica* and *japonica*, presumably due to intensive genetic admixture by breeding over history (Zhao et al, 2010; Xu et al, 2012). However, the variability given multiple traits is substantially distinguishable between *indica* and *japonica* because multiple layers of the narrowly distinguishable effects collectively magnify differences between *indica* and *japonica*. The highest AUC obtained by the LRM accounts for 0.9977 (Table 1) in a 6D selection including the predictors number per plant, seed number per panicle, florets per panicle, panicle fertility, straighthead susceptibility and blast resistance. Importantly, the panicle-related traits, straighthead susceptibility and blast resistance frequently appeared across all sets. These findings may be related to previous knowledge that the variability in panicle characteristics, straighthead susceptibility and blast resistance is strongly associated with the *indica* and *japonica* classification; panicles are long and sparse in *indica* but short and dense in *japonica* (Bai et al, 2016); straighthead resistance and blast resistance are greater in *indica* than *japonica* (Yan et al, 2005; Jia et al, 2011). Though I have presented only the best selection of predictors in each set, other selections of predictors producing a certain AUC level or greater (e.g. 0.9) will also provide useful insight into which traits collectively characterize the *indica/japonica* classification. The dendrogram-based method using 36,901 SNP variables was perfectly effective with an AUC = 1 (Figure 2). Considering that the LRM-based method achieved an AUC = 0.9977 with only six predictors, its performance is still highly impressive.

The ultimate purpose of this study is to demonstrate the effectiveness of the LRM-based *indica/japonica* predictions of unclassified Asian rice cultivars. To estimate the *indica/japonica* prediction accuracy, I conducted two exams using Equation 8: the first exam was to conduct the *indica/japonica* predictions in each single minor subpopulation (*admixed, aromatic, aus*); and the second exam was to perform the 100 times iterative, ten-fold cross-validations given the *indica/japonica* accessions. As a result of the first exam, the agreements between two results obtained by (1) the LRM-based method and (2) the dendrogram-based method varied, which account for 92.3 % (48/52) in the *admixed* group, 66.7% (8/12) in the *aromatic* group and 76.9 % (20/26) in the *aus* group (Figure 4). According to the previous knowledge, *aus* is related to *indica*, and *aromatic* is in between *indica* and *japonica* (Garris et al, 2005; Zhao et al, 2011; McCouch et al, 2016; Chin et al, 2017). From this perspective, the result obtained by the dendrogram-based method seemed to be perfectly accurate in that the *aus* and *indica* groups are separately clustered in the *indica* clade, the *aromatic* group is distantly related to the *japonica* group in the *japonica* clade, and the *admixed* accessions are spread across all subpopulation groups (Supplementary figure 2). The *indica/japonica* prediction accuracy of the LRM can be improved by calculating parameters in a condition where the *aromatic* and *aus* groups are assigned to the *japonica* and *indica*, respectively. As a result of the second exam, the *indica/japonica* prediction accuracy resulting from 100 times ten-fold CVs accounted for 0.9779. This result shows that, given pure *indica* and *japonoica* accessions, the *indica/japonica* prediction of the LRM is highly effective.

## Conclusion

In Asian cultivated rice, the use of the LRM will provide rice breeders with a precise and fast *indica/japonica* classification tool promising in-depth insights into the *indica/japonica* classification by phenotype.

